# Revealing subthreshold motor contributions to perceptual confidence

**DOI:** 10.1101/330605

**Authors:** Thibault Gajdos, Stephen M. Fleming, Marta Saez Garcia, Gabriel Weindel, Karen Davranche

## Abstract

Established models of perceptual metacognition, the ability to evaluate our perceptual judgments, posit that perceptual confidence depends on the strength or quality of feedforward sensory evidence. However, alternative theoretical accounts suggest the entire perception-action cycle, and not only variation in sensory evidence, is monitored when evaluating confidence in one’s percepts. Such models lead to the counterintuitive prediction that perceptual confidence should be directly modulated by features of motor output. To evaluate this proposal here we recorded electromyographic (EMG) activity of motor effectors while subjects performed a near-threshold perceptual discrimination task and reported their confidence in each response. A subset of trials exhibited sub-threshold EMG activity in response effectors before a decision was made. Strikingly, trial-by-trial analysis showed that confidence, but not accuracy, was significantly higher on trials with subthreshold motor activation. These findings support a hypothesis that preparatory motor activity impacts upon confidence over and above performance, consistent with models in which perceptual metacognition integrates information across the perception-action cycle.

## 1. Introduction

Our perception of the outside world is associated with variable degrees of confidence. For instance, when driving in fog, we are less confident about the presence of oncoming cars and may slow down accordingly. Prominent computational models, grounded in signal detection theory and evidence accumulation frameworks, suggest that perceptual confidence is determined by the strength of an internal decision variable that encodes evidence in support of different interpretations of a stimulus (Vickers, 1979; Galvin, 2003; Kiani & Shadlen, 2009). Intuitively, the stronger the perceptual evidence, the more confident we should be in our decision. Such models in turn predict that perceptual discrimination accuracy and confidence should be tightly coupled, and be affected by similar experimental manipulations.

However, recent theoretical models of metacognition suggest that the entire perception-action cycle, and not only variation in sensory evidence, may be monitored when evaluating confidence in one’s decisions (Pouget et al., 2016; Fleming & Daw, 2017). Such models lead to the counterintuitive prediction that confidence in perceptual judgments – but not perceptual discrimination accuracy – should be modulated by features of motor output as well as perceptual input. Indeed, recent observations suggest that confidence in perceptual tasks does not only result from feedforward sensory input, but also depends on other sources of information (Siedlecka et al., 2016; 2018). For instance longer response times (Kiani, Corthell & Shadlen, 2014) or unexpected arousal (Allen et al., 2016) have been found to modulate confidence without affecting accuracy. Confidence in perceptual tasks is also disrupted by manipulation of movement speed (Palser, Fotopoulou & Kilner, 2018) and transcranial magnetic stimulation of the motor system (Fleming & al., 2015). However, a link between trial-by-trial motor output and perceptual confidence remains to be established.

Here we set out to study this relationship by recording subthreshold fluctuations in electromyographic (EMG) activity as sensitive markers of motor preparatory activity during perceptual discrimination. Specifically, partial muscular activations (henceforth, partial activations) corresponding to sub-threshold motor responses have repeatedly been observed in between-hand choice reaction time tasks (Eriksen, Coles, Morris & O’hara, 1985; Hasbroucq, Possamaï, Bonnet & Vidal, 1999; Meckler, Carbonnell, Ramdani, Hasbroucq & Vidal, 2017). These partial activations are typically followed by an error-related negativity (Ne) in the EEG signal (Scheffers, Coles, Bernstein, Gehring, & Donchin, 1996; Vidal, Hasbroucq, Grapperon, & Bonnet, 2000; Meckler, Carbonnell, Ramdani, Hasbroucq & Vidal, 2017), and intracerebral recordings demonstrate that the Ne is elicited in supplementary motor area (SMA) and pre-SMA / rostral cingulate (Bonini, Burle, Liégeois-Chauvel, Régis, Chauvel & Vidal, 2014), thereby establishing a direct link between a component of motor output (partial activation) and frontal lobe structures involved in performance monitoring and metacognition (Carter, Braver, Barch, Botvinick, Noll & Cohen, 1998; Coull, Vidal & Burle, 2016; Fleming, van der Putten & Daw, 2018). Notably, variations in error-related EEG potentials have also been recently linked to subjective confidence in perceptual decisions (Boldt & Yeung, 2015). Taken together, these findings suggest a hypothesis in which partial activations are specifically associated with modulation of retrospective confidence but not task performance (there is no prior evidence that partial activations impact discrimination accuracy).

Here we recorded EMG activity of response agonists while participants performed a difficult perceptual discrimination task by pressing the appropriate key with the left or right thumb. After each trial, they were required to verbally provide their confidence in their response. Trial-by-trial analysis showed that confidence was significantly higher on trials with partial activation, while no significant effect was observed on accuracy. These findings support a hypothesis that subthreshold motor output impacts upon confidence over and above performance, and are consistent with models in which confidence integrates information across the perception-action cycle.

## 2. Results

To test whether partial activations modulate confidence, we analysed data from 19 participants (see Methods for details) who performed a perceptual discrimination task described in Figure 1. On each trial, a low-contrast grating was presented centrally for 33 ms. Participants were instructed to indicate, as quickly as possible, whether the grating was oriented vertically or horizontally by pressing the appropriate response key with the right or the left thumb. During each trial, EMG activity of the *flexor pollicis brevis* of each thumb and force production were recorded. After making their decision, subjects were asked to rate their confidence verbally on a scale from 1 (low confidence) to 4 (high confidence). Each subject performed 700 trials evenly split into ten blocks separated by self-paced rest periods. During a preliminary calibration block, stimulus contrast was manipulated with a 1-up-3-down staircase procedure expected to yield 80% correct responses. On each trial, participants were also allowed to report if they consciously detected making an error. These trials (2% of the total number of trials) were omitted from further analysis.

**Figure 1.**
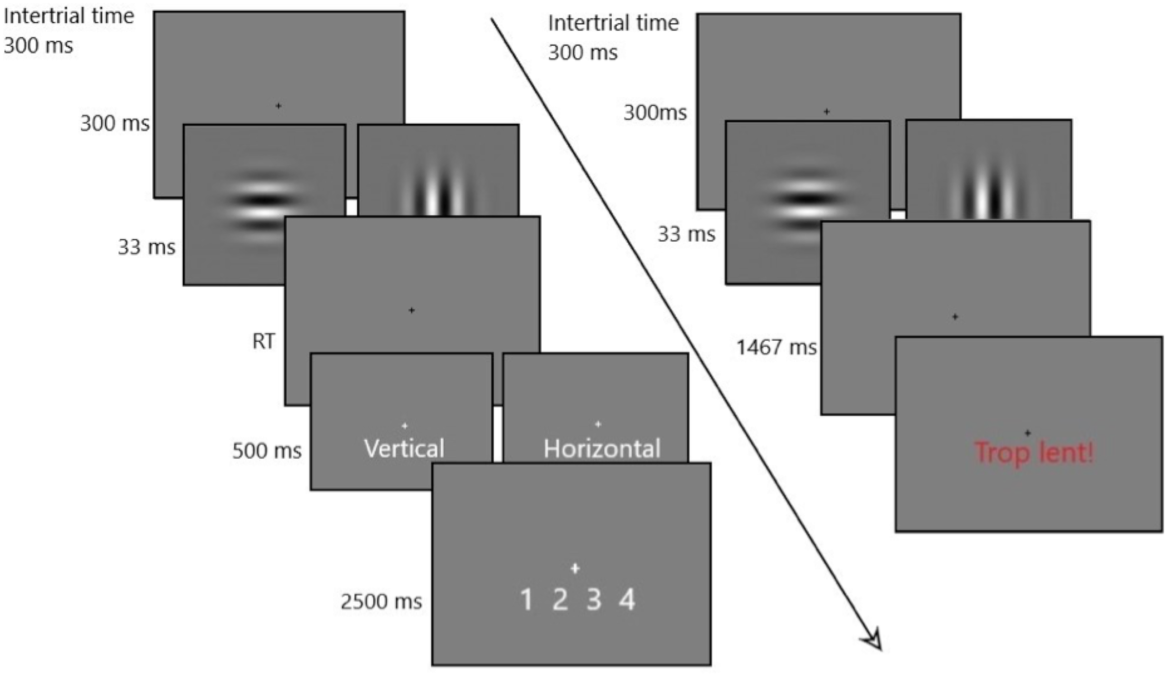
Trial sequence. Successfully completed trial (left). A fixation point was displayed for 300 ms. A Gabor patch (shown in the image) briefly appeared (33 ms) followed by another fixation point. Subjects were required to respond as quickly and accurately as possible according to whether the stimulus was oriented vertically or horizontally by pressing the appropriate key in less than 1500 ms. Following the response an answer confirmation was displayed for 500 ms. Subjects then had 2500 ms to verbally rate their confidence on a scale from 1 (low confidence) to 4 (high confidence). Failed trial (right). If a response was not provided in less than 1500 ms, the French words “Trop lent!” (“too slow”) were displayed, and the next trial began after 300 ms.

EMG traces were inspected off-line, trial by trial, as displayed on a computer screen and the onsets of the changes in activity were hand-scored. After visual inspection, 4% of the trials were rejected because of tonic activity or artefacts. Trials with more than one partial activation were discarded from further analysis. Remaining trials (*11471, mean* = 604 per subject, *SD* = 64) were classified into three categories: trials without partial activation (*mean* = 83%, *SD* = .09), trials with a partial activation ipsilateral to the provided response (mean = 8.5%, *SD* = .05), and trials with a partial activation contralateral to the provided response (mean = 8.3%, *SD* = .06). The average time course of EMG for trials with and without partial activation is shown in Figure 2. Reaction time was measured between the onset of the stimulus and the onset of the required motor response. As expected, partial activations were more frequent among the slowest trials, on which there is more time for them to occur. Mean accuracy for the orientation discrimination was 80% (*SD* = .11), and mean confidence was 2.6 (*SD* =.32).

**Figure 2:**
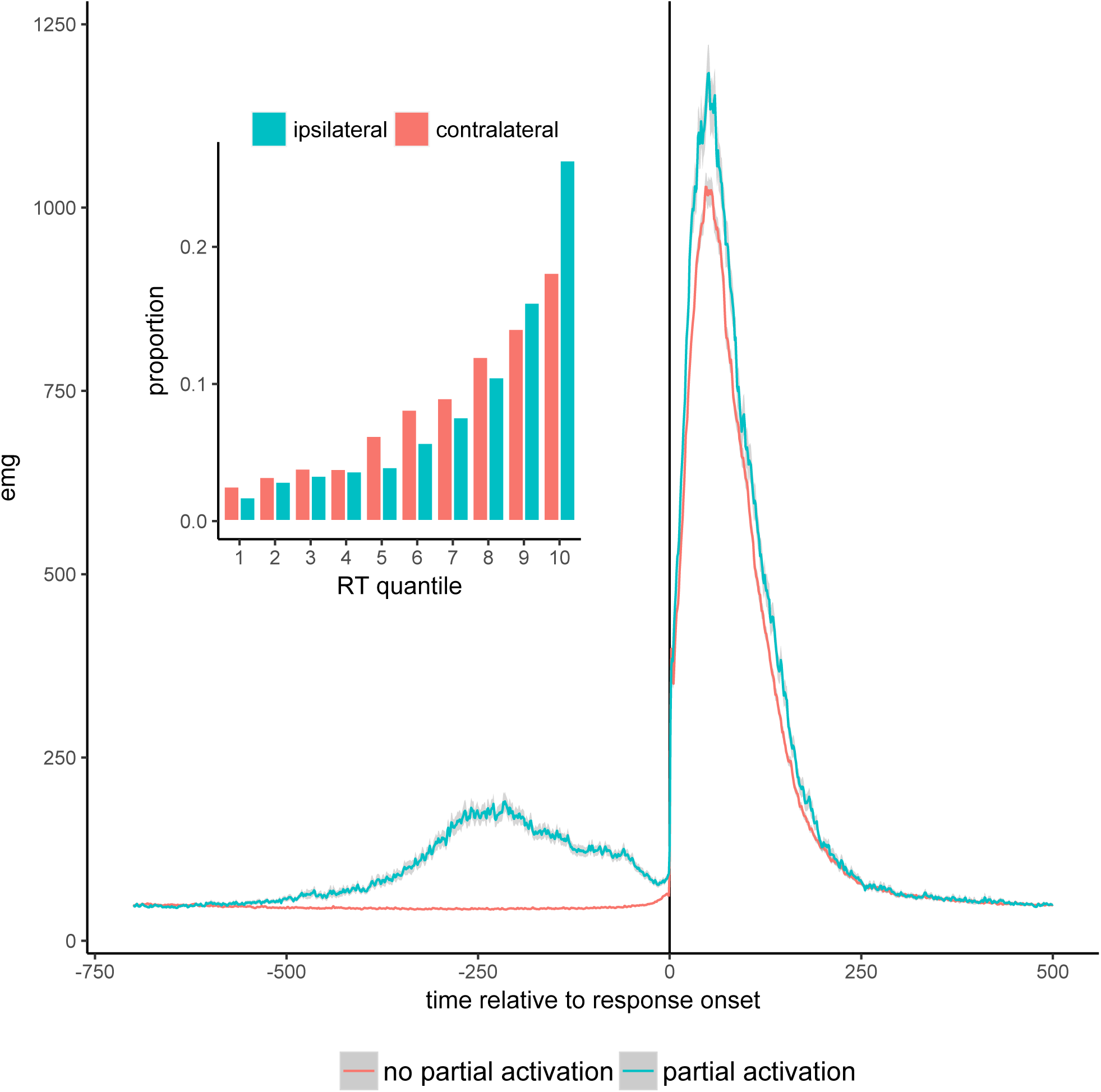
Main figure: Average EMG time course for trials with (blue) and without (red) partial activation. Shaded areas represent 95% confidence. Trials were aligned to response onset (time = 0). Inset figure shows proportion of partial activation (ispsilateral (blue) and contralateral (red) to the response) by RT quantile, averaged across subjects.

### 2.1. Confidence and accuracy

We analysed the effects of partial activation on confidence within a hierarchical linear mixed-effects model, including the intercept, partial activation (separately according to whether it is ipsilateral or contralateral to the response), response accuracy, force production and reaction time as fixed and random effects (see Methods for details). Results of the regression analysis are reported in Table 1. As expected, confidence was higher for correct vs. error trials (β= .60, *SE* = .055, *p* < .001) and faster vs. slower responses (β = −2.3, *SE* = .18, *p* < .001) (Henmon, 1911; Baranski & Petrusic, 1998; Kiani et al., 2014). Crucially, there was also a significant effect of partial activation on confidence: participants reported higher confidence for trials with compared to without partial activation. Interestingly, this effect was similar for ipsilateral (β = .15, *SE* = .040, *p* = .002) and contralateral (β = .16, *SE* = .049, *p* = .004) partial activations (effect size of approximately 0.15 confidence SD). Thus, while partial activations were associated with a significant effect on confidence, the congruency between partial activation and response had no effect on confidence. Finally, confidence was also higher when responses were provided with stronger force (β = .33, *SE* =.11, *p* = .007).

**Table 1:** Hierarchical regression coefficients predicting confidence from accuracy, ispilateral and contralateral partial activations, reaction time and force production. Predictors were coded as follows – Accuracy: error = 0, correct = 1; Ipsilateral: absent = 0, present = 1; Contralateral: absent = 0, present = 1. ^**\***^ p < .05, ^**\*\***^ p < .01, ^***^ p < .001. Reaction time and force production are median- centered

| Predictor | $\beta$ | $p$ |
| --- | --- | --- |
| Intercept | 2.1*** (.12) | < .001 |
| Accuracy | .60*** (.055) | < .001 |
| Reaction time | -2.3*** (.18) | < .001 |
| Force production | .33** (.11) | .007 |
| Ipsilateral | .15** (.040) | .002 |
| Contralateral | .16** (.049) | .004 |

Number of subjects: 19. Number of observations: 11471

To further understand the drivers of the partial activation effect on confidence, we visualized confidence and accuracy data as a function of partial activation and response time (Figure 3). The lower panel of Figure 3 illustrates an overall decrease of confidence as RT increases, together with a consistent boost in confidence within a majority of RT quantiles on trials with partial activations. Notably however there was no consistent effect of partial activation on first-order performance (Figure 3, upper panel). A lack of effect of partial activation on accuracy was confirmed within a generalized hierarchical linear mixed-effects model, including the intercept, partial activation (ipsilateral or contralateral) and their interactions, as well as force production and reaction time as fixed and random predictors. Results of the regression analysis are reported in Table 2. Accuracy was higher for faster responses (β = - 2.3, *SE* = .40, *p*<.001). Importantly, however, partial activations, either ipsilateral or contralateral, were unrelated to accuracy on a trial by trial basis (all *p* s>.35). Force production also had no significant effect on accuracy (*p* = .49).

**Table 2:** Hierarchical regression coefficients predicting accuracy from ispilateral and contralateral partial activations, Reaction time and force production. Predictors were coded as follows – Ipsilateral: absent = 0, present = 1; Contralateral: absent = 0, present = 1. ^**\***^p < .05, ^**\*\***^p < .01, ^***^p<.001. Number of subjects: 19. Number of observations: 11471

| Predictor | $\beta$ | p |
| --- | --- | --- |
| Intercept | 1.8*** (.22) | <.001 |
| Reaction time | -2.3*** (.40) | < .001 |
| Force production | .13 (.19) | .49 |
| Ipsilateral | -.11 (.15) | .44 |
| Contralateral | .14 (.15) | .35 |

**Figure 3:**
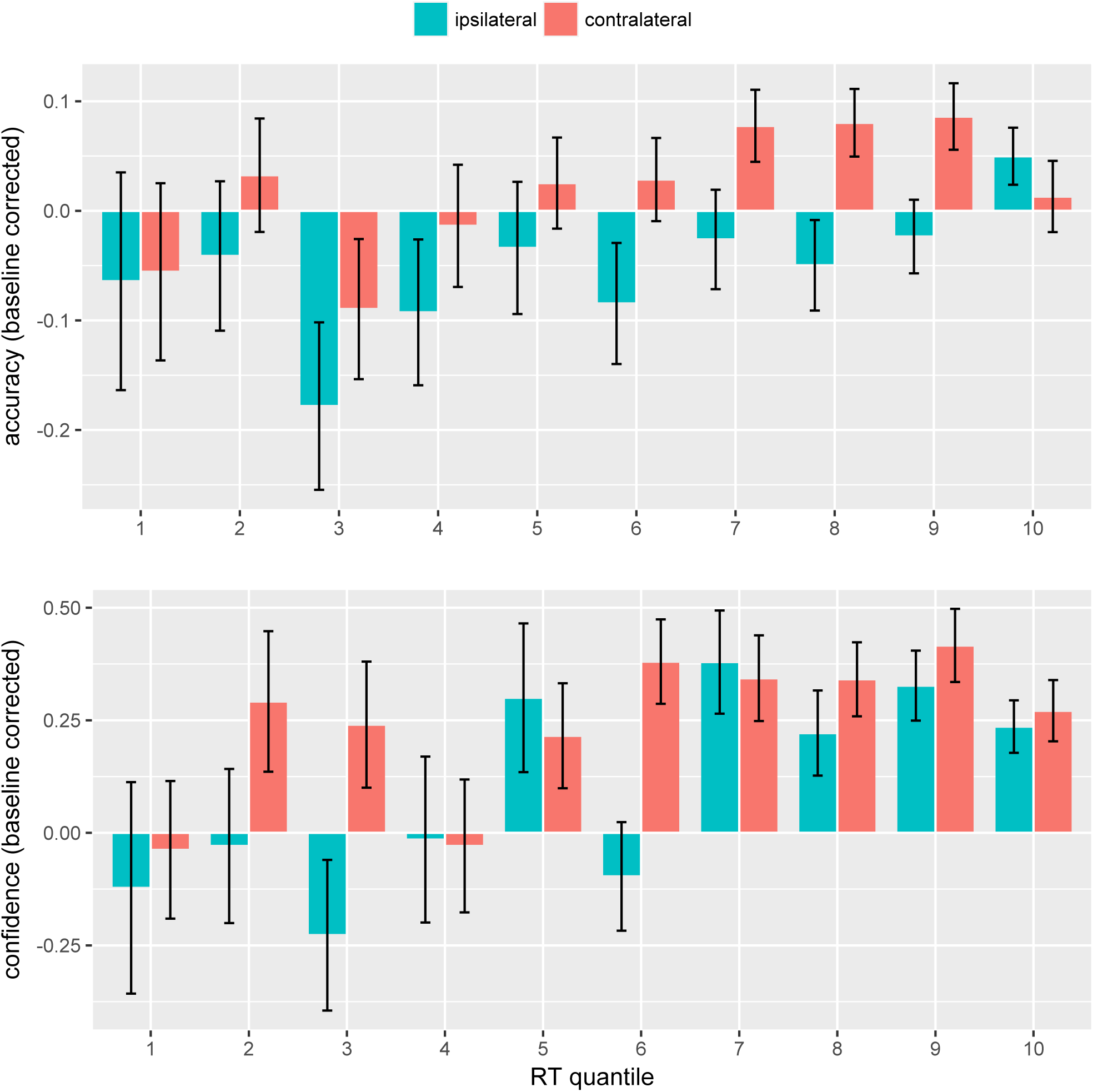
Mean accuracy (top panel) and confidence (bottom panel) by RT quantiles across subjects for trials without partial activation (green), and with partial activation ispsilateral (blue) or contralateral (red) to the response, baseline-corrected with respect to trials without partial activation. Error bars reflect standard errors of the mean.

Interestingly, inspection of Figure 3 indicated that the effect of partial activations on confidence effect was most prominent for the slowest RT quantiles, suggesting an interaction between RT and partial activation. To formally test for such an effect, we replicated the previous analysis, now including an interaction term between RT and partial activation (complete results are presented in Supplementary Material, table S1.1). We indeed found a significant interaction between RT and contralateral partial activations (β = .40, *SE* = .17, *p* = .004), with a similar trend for ipsilateral activations (β = .50, *SE* = .26, *p* = .07). An interaction between RT and partial activations might be explained by fewer partial activations among slow trials, resulting in a lack of sufficient statistical power to detect their potential effect on confidence. Alternatively, it might be partial activations become a cue to confidence specifically on longer trials (see Figure S1.1 in Supplementary Material). The current data do not allow us to decide between these two explanations.

It has been shown that faster reaction times are used as a cue to boost confidence (Kiani et al., 2014). One might therefore wonder whether the impact of partial activations on confidence is a secondary result of an early motor activation being interpreted as a marker of a fast response during confidence formation. Under this explanation, we should observe an impact of the timing of partial activations on confidence, with earlier activations corresponding to higher confidence levels.

To test this hypothesis, we repeated our previous analysis, adding the onset time of the first muscular activation (absolute pre-motor time, Apmt) as a predictor in the regression. Note that in the absence of partial activation, Apmt corresponds to the earliest detectable time of the actual response. In opposition to an explanation based on response timing, shorter Apmt values had no significant impact on confidence level (β *=* -.19, *SE* = .22, *p* = .39). Therefore, the influence of partial activation on confidence cannot be explained by the fact that they occur earlier than the final response (see Supplementary Material, Table S1.2).

## 3. Discussion

Our results reveal that partial muscular activation of response effectors has a specific impact on perceptual confidence, without altering primary perceptual discrimination performance. More precisely, we found that partial activations systematically boost confidence, without impacting accuracy.

At first glance, such “early” effects of motor activation may indicate that they influence metacognitive evaluation in the same manner as faster responses, acting as a cue to higher confidence. Indeed, it has been shown that response times directly affect confidence levels in perceptual discrimination (Kiani & al., 2014). However, in contradistinction to this hypothesis, we did not find any influence of the timing of partial activation on confidence level, despite finding a significant effect of overall response time. This suggests that partial activation represents a distinct influence on confidence, rather than another marker of faster responding. Notably, these effects of partial activation on confidence were obtained in the absence of a similar changes in accuracy. If anything, ipsilateral activations *reduced* accuracy, while still boosting confidence (compare RT bins 7, 8 and 9 in Figure 3). In a regression model of accuracy that included response times, we found a significant positive interaction of response time with ipsilateral (β = 1.2, *SE* = .47, *p* = .009), but not contralateral (β = -.32, *SE* = .62, *p* = .61) partial activations, together with a significant negative main effect (β = - .40, *SE* = .16, *p*<.01) (see table S1.3 and Figure S1.2 in the Supplementary Material). These findings suggest that any effect of partial activation on first-order performance is qualitatively different from its effect on confidence.

Our results build on and extend previous studies suggesting that the motor system itself contributes to metacognitive judgments in decision-making (Palser & al., 2018; Fleming & al., 2015; Siedlecka & al., 2016, 2018). However, our study represents the first to demonstrate a specific impact of subthreshold motor activation – as measured using EMG – on perceptual confidence, and therefore provides a particularly strong and direct test of this hypothesis. Moreover, the impact of partial activation on confidence was independent of the congruence with the provided response: ispilateral and contralateral partial activations have a similar effect on confidence. At first glance this might seem surprising, as it has been previously shown that single-pulse transcranial magnetic stimulation (TMS) increases confidence when applied to the hemisphere associated to the chosen response, but decreases it when applied to the hemisphere associated to the alternative response (Fleming & al., 2015). Further research, potentially combining motor TMS with measurement of partial activation, is needed to understand the relation between these effects. One potential hypothesis is that partial motor activations and pre-motor activity affect confidence judgements through distinct mechanisms. It has been shown that partial activations are followed by a Ne (Meckler, Carbonnell, Ramdani, Hasbroucq & Vidal, 2017) originating in SMA proper (Bonini & al., 2014), a structure involved in performance monitoring (Coull & al. 2016). It might thus be the case that partial activations are interpreted as efficiently corrected errors, leading to higher confidence in the final decision. By contrast, stimulation of lateralised pre-motor cortex is more likely to directly influence the representations associated with a particular response, and therefore influence confidence in an asymmetric manner. Our data do not provide the means to directly test this explanation, which we leave open for future research.

In summary, our findings contribute to growing evidence that confidence in perceptual decisions not only depends on feedforward sensory evidence, but also depends on motor information. They also suggest that different components of motor output might contribute differently to confidence. Together our findings are broadly consistent with “second-order” models in which metacognitive confidence integrates information across the perception-action cycle (Fleming & Daw, 2017). Much remains to be done to understand how exactly these components are combined and processed into metacognitive judgments.

## 4. Methods and Materials

### 4.1. Participants

Sample size was based on previous published study investigating the role of pre-motor activation on confidence (Fleming et al., 2015). We decided prior to data collection to ensure data analysis would be possible for between 16 and 25 participants. A total of 27 participants with normal or corrected to normal vision and no history of neurological or psychiatric disorders thus participated in the study. Five participants were excluded before analysis either because the EMG signal was too noisy (2 participants), they did not comply with the instructions (2 participants), or they failed to accomplish the task (1 participant). Furthermore, three outliers were excluded because they displayed multiple activations in more than 70% of the trials (2 participants), or used the highest level of confidence in more than 80% of the trials (1 participant). We thus analysed data of 19 participants in the final sample (11 females, 7 males; mean age = 23 years, SD 3.7). We note that results are qualitatively similar if outliers are not excluded (see Appendix 2).

### 4.2. Stimuli and procedure

The experiment was carried out at the Laboratoire de Neurosciences Cognitives, in Marseille (France). Participants performed the experiment in a dark and sound-shielded Faraday cage. They were seated 100 cm in front of a 15-inch CRT monitor with a refresh rate of 60 Hz. Response buttons were be fixed on the top of two plastic cylinders (diameter 3 cm, height 7.5 cm), 20 cm apart. Participants responded by exerting at least 600 mg of pressure on a button. Responses were confirmed by audio feedback.

Stimuli were generated using the Psychopy library for Python (Peirce, 2009). Each trial started with a fixation cross, displayed for 300 ms. It was followed by a low-contrast grating randomly oriented either horizontally or vertically, and presented centrally for 33 ms (4° diameter). Participants were instructed to indicate, as quickly as possible, whether the grating was vertically or horizontally oriented by pressing the appropriate response key with their right or left thumb. The association between grating orientation (horizontal or vertical) and response key (left or right) was counterbalanced across subjects. Responses faster than 1500 ms were confirmed by a message reporting the provided answer (horizontal or vertical) displayed for 500 ms. Responses longer than 1500 ms were omitted from analysis and feedback was provided to the subject signalling that her answer was too slow. After confirmation of the response, subjects were asked to loudly rate their confidence from 1 (low confidence) to 4 (high confidence) while a scale ranging from 1 to 4 was displayed on the screen for 2500 ms. If they consciously detected making a mistake they were asked to say “error” instead of their confidence level. Responses were recorded and written down by the experimenter, who was outside the room where the participants sat. Each trial lasted 4,333 ms.

For each participant, stimulus contrast was individually adjusted in a preliminary session using a 1- up-3-down staircase procedure. The experiment consisted of 699 trials, split into 9 blocks of 70 trials each and one block of 69 trials. Rest periods between each block were self-paced.

### 4.3. EMG and force recording

Electromyographic (EMG) activitiy was recorded continuously from preamplified Ag/AgCl electrodes (Biosemi® Active-Two electrodes®, Amsterdam), pasted onto the skin of the thenar eminence over the *flexor pollicis brevis* of each thumb, about two centimetres apart. The common reference electrode was situated above the first vertebra. The signal was filtered and digitized on-line (bandwidth: 0–268 Hz, 3 dB/octave, sampling rate: 1024 Hz). The experimenter continuously monitored the signal and asked subjects to relax their muscles if the signal became noisy.

The thumb force production was measured as a force signal and digitized on line (A/D rate 2 kHz) allowing us to record the force applied by the participant and trigger a response signal when a force threshold of 600g is exceeded.

### 4.4. Data analysis

No statistical analyses were conducted prior to having completed the collection of data for all participants. The protocol was registered on Open Science Framework before starting the experiment (Saez, Gajdos, Fleming & Davranche, 2017).

#### 4.4.1. EMG analysis

The recorded EMG signals were analysed off-line. The trace corresponding to the EMG was displayed on the computer screen and the EMG onsets were hand-scored. Importantly, at this stage the experimenter was unaware of the confidence scores registered separately, avoiding any bias in analysis. Reaction time was measured between the onset of the stimulus and the required motor response. We also defined Absolute Pre-Motor Time (Apmt) as the delay between the onset of the stimuli and the beginning of the first muscular activation.

#### 4.4.2. Trials exclusions

Force production was not registered for 50 trials (.4% of the total number of trials), due to the force peak latency occurring more than 1500 ms following stimulus onset. After visual inspection, 715 trials (5.4% of the total number of trials) were discarded from further analysis due to the presence of tonic activity or artefacts. Trials reported as errors by subjects were also disregarded (442 trials, 3.3% of the total number of trials). Finally, trials with more than one partial activation were excluded in initial analyses (779 trials, 5.9% of the total number of trials), but were included in the regression analysis reported in Table 4. In our primary analyses we excluded a total of 1810 trials (13.6%), and analysed 11471 trials (mean = 604 per subject, *SD* = 64).

#### 4.4.3. Statistical procedure

The influence of partial activation on confidence and accuracy was analysed with hierarchical linear and generalized mixed-effects models, respectively, using the lmer4 package (Bates, Mächler, Bolker, & Walker, 2015) in R (version 4.2, R Core Team, 2017). All regressions were performed with the restricted maximum likelihood fitting method, and *p* values for coefficients were obtained with the lmerTest package (Kuznetsova, Brockhoff, & Christensen, 2017). We used an alpha level of .05 for all statistical tests.

Predictors were coded using treatment contrasts (thus “accuracy” takes a value of 1 if the trial is correct, and 0 otherwise, “ispilateral” takes a value of 1 if there is a partial ipsilateral activation, and 0 otherwise; “contralateral” takes value of 1 if there is a partial contralateral activation, and 0 otherwise). Reaction time and Absolute Pre-Motor Time (Apmt) were measured in seconds. Force was measured in kilograms. Reaction time, Absolute Pre-Motor Time and force are median-centered. To keep models tractable, we assumed zero correlation between random effects.

We checked that results were robust to the inclusion of outliers (Appendix 2) and we replicated all analyses using Bayesian methods as implemented in the Brms package in R (Appendix 3) (Bürkner, 2016).

## Appendix 1: Supplementassry material

**Table S1.1:** Hierarchical regression coefficients predicting confidence from accuracy, ispilateral and contralateral partial activations, reaction time and force production. Predictors were coded as follows – Accuracy: error = 0, correct = 1; Ipsilateral: absent = 0, present = 1; Contralateral: absent = 0, present = 1. ^**\***^p < .05, ^**\*\***^p < .01, ^***^p< .001. Number of subjects: 19. Number of observations: 1147

| Predictor | $\beta$ | p |
| --- | --- | --- |
| Intercept | 2.1*** (.12) | < .001 |
| Accuracy | .59*** (.055) | < .001 |
| Reaction time | -2.4*** (.18) | < .001 |
| Force production | .32** (.10) | .007 |
| Ipsilateral | .071 (.044) | .13 |
| Contralateral | .12* (.050) | .026 |
| Reaction time x Ipsilateral | .50 (.26) | .071 |
| Reaction time x Contralateral | .49** (.17) | .004 |

**Figure S1.1:**
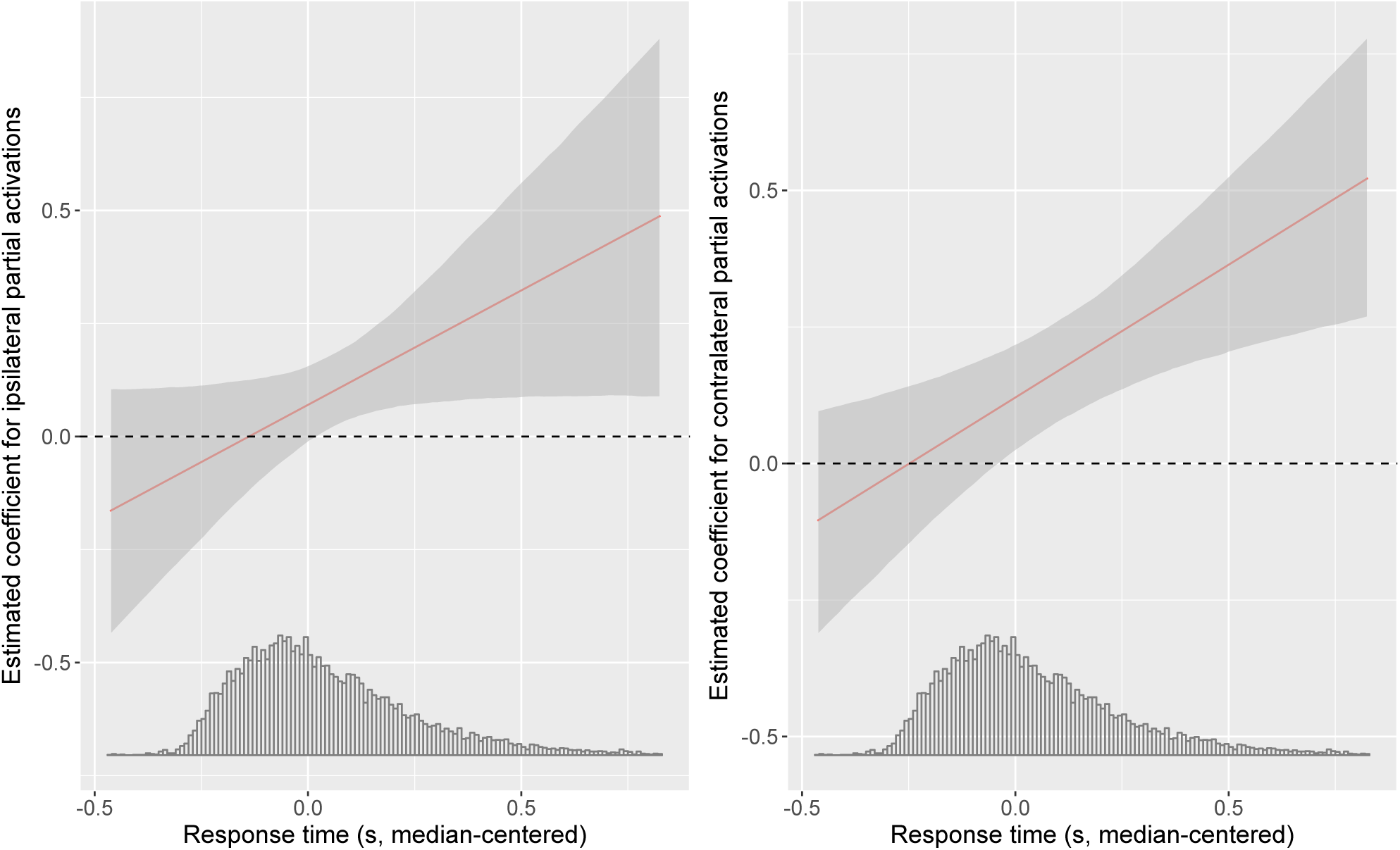
Visualisation of interaction between RT and partial activation. Estimated coefficient for the impact of ipsilateral (left panel) and contralateral (right panel) partial activations on confidence. Shaded region represents 95% confidence. Inset figures show the distributions of ipsilateral and contralateral partial activations onset times.

**Table S1.2:** Hierarchical regression coefficients predicting confidence from accuracy, ispilateral and contralateral partial activations, absolute pre-motor time (Apmt) and force production. Predictors were coded as follows – Accuracy: error = 0, correct = 1; Ipsilateral: absent = 0, present = 1; Contralateral: absent = 0, present = 1. ^**\***^p < .05, ^**\*\***^p < .01, ^***^p< .001. Number of subjects: 19. Number of observations: 11471

| Predictor | $\beta$ | p |
| --- | --- | --- |
| Intercept | 2.1*** (.12) | < .001 |
| Accuracy | .59*** (.055) | < .001 |
| Reaction time | -2.2*** (.23) | < .001 |
| Apmt | - .19 (.22) | .39 |
| Force production | .33** (.11) | .007 |
| Ipsilateral | .12** (.045) | .009 |
| Contralateral | .14* (.055) | .02 |

**Table S1.3:** Hierarchical regression coefficients predicting accuracy from ispilateral and contralateral partial activations, reaction time and their interactions, and force production. Predictors were coded as follows – Accuracy: error = 0, correct = 1; Ipsilateral: absent = 0, present = 1; Contralateral: absent = 0, present = 1. ^**\***^p < .05, ^**\*\***^p < .01, ^***^p< .001. Number of subjects: 19. Number of observations: 1147

| Predictor | $\beta$ | p |
| --- | --- | --- |
| Intercept | 1.8*** (.22) | < .001 |
| Reaction time | -2.4*** (.41) | < .001 |
| Force production | .12 (.19) | .52 |
| Ipsilateral | -.40* (.16) | .011 |
| Contralateral | .22 (.14) | .13 |
| Reaction time x Ipsilateral | 1.2** (.47) | .009 |
| Reaction time x Contralateral | -.32 (.62) | .61 |

**Figure S1.2:**
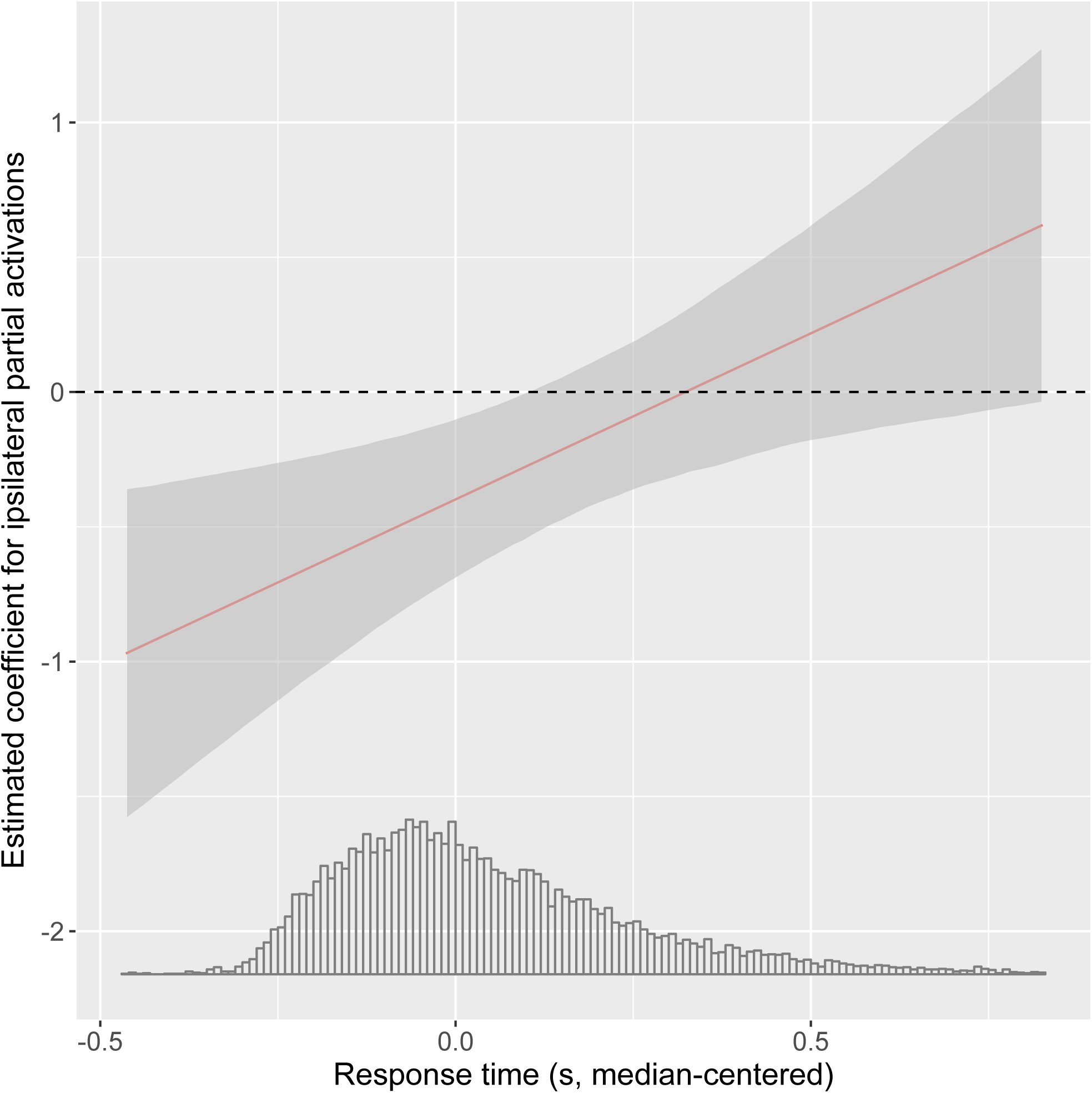
Visualisation of interaction between RT and ipsilateral activation. Estimated coefficient for the impact of ipsilateral partial activations on accuracy. Shaded region represents 95% confidence. Inset figure show the distribution of ipsilateral partial activations onset times.

## Appendix 2: Replication without excluding outliers

**Table S2.1:** Predicting Confidence from Accuracy, Ispilateral and Contralateral partial activation, Reaction time and Force production. Predictors were coded as follows – Accuracy: error = 0, correct = 1; Ipsilateral: absent = 0, present = 1; Contralateral: absent = 0, present = 1. ^**\***^p < .05, ^**\*\***^p < .01, ^***^p< .001. Number of subjects: 22. Number of observations: 12645

| Predictor | $\beta$ | p |
| --- | --- | --- |
| Intercept | 2.1<sup>***</sup> (.13) | < 0.001 |
| Accuracy | .71<sup>***</sup> (.095) | < 0.001 |
| Reaction time | -2.2<sup>***</sup> (.16) | < 0.001 |
| Force production | .35<sup>**</sup> (.096) | .002 |
| Ipsilateral | .11<sup>**</sup> (.039) | .009 |
| Contralateral | .13<sup>**</sup> (.044) | .009 |

**Table S2.2:** Predicting Confidence from Accuracy, Ispilateral and Contralateral partial activation, Reaction time and Force production. Predictors were coded as follows – Accuracy: error = 0, correct = 1; Ipsilateral: absent = 0, present = 1; Contralateral: absent = 0, present = 1. ^**\***^p < .05, ^**\*\***^p < .01, ^***^p< .001. Number of subjects: 22. Number of observations: 12645

| Predictor | $\beta$ | p |
| --- | --- | --- |
| Intercept | 2.1<sup>***</sup> (.13) | < 0.001 |
| Accuracy | .70<sup>***</sup> (.095) | < 0.001 |
| Reaction time | -2.3<sup>***</sup> (.17) | < 0.001 |
| Force production | .34<sup>**</sup> (.093) | .002 |
| Ipsilateral | .040 (.040) | .33 |
| Contralateral | .097<sup>*</sup> (.044) | .04 |
| Reaction time x Ipsilateral | .54<sup>*</sup> (.22) | .02 |
| Reaction time x Contralateral | .49<sup>**</sup> (.15) | .001 |

**Table S2.3:**
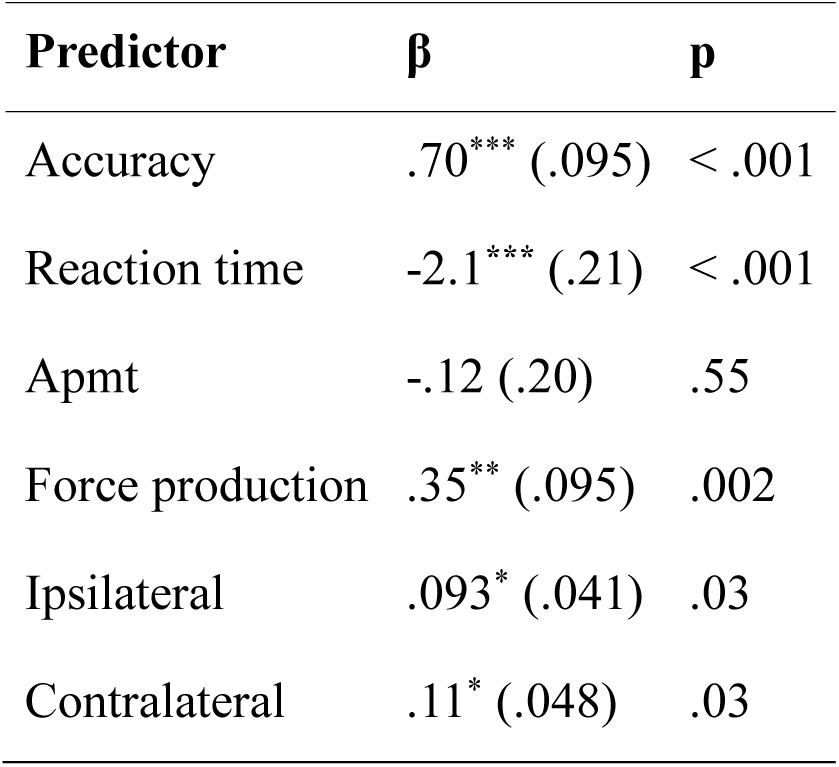
Hierarchical regression coefficients predicting confidence from accuracy, ispilateral and contralateral partial activations, absolute pre-motor time (Apmt) and force production. Predictors were coded as follows – Accuracy: error = 0, correct = 1; Ipsilateral: absent = 0, present = 1; Contralateral: absent = 0, present = 1. ^**\***^p < .05, ^**\*\***^p < .01, ^***^p< .001. Number of subjects: 22. Number of observations: 12645

**Table S2.4:** Hierarchical regression coefficients predicting accuracy from ispilateral and contralateral partial activations, Reaction time and force production. Predictors were coded as follows – Ipsilateral: absent = 0, present = 1; Contralateral: absent = 0, present = 1. ^**\***^p < .05, ^**\*\***^p < .01, ^***^p< .001. Number of subjects: 22. Number of observations: 12645

| Predictor | $\beta$ | p |
| --- | --- | --- |
| Intercept | 1.7*** (.21) | <.001 |
| Reaction time | -2.2*** (.39) | < .001 |
| Force production | .29 (.19) | .13 |
| Ipsilateral | -.17 (.13) | .20 |
| Contralateral | .21 (.14) | .12 |

**Table S2.5:** Hierarchical regression coefficients predicting accuracy from ispilateral and contralateral partial activations, Reaction time and force production. Predictors were coded as follows – Ipsilateral: absent = 0, present = 1; Contralateral: absent = 0, present = 1. ^**\***^p < .05, ^**\*\***^p < .01, ^***^p< .001. Number of subjects: 22. Number of observations: 12645

| Predictor | $\beta$ | p |
| --- | --- | --- |
| Intercept | 1.7*** (.21) | <.001 |
| Reaction time | -2.3*** (.40) | < .001 |
| Force production | .28 (.19) | .14 |
| Ipsilateral | -.40** (.13) | .003 |
| Contralateral | .27* (.12) | .026 |
| Reaction time x Ipsilateral | 1.1** (.40) | .006 |
| Reaction time x Contralateral | -.39 (.58) | .50 |

## Appendix 3: Bayesian analyses

Bayesian analyses were implemented using the Brms package in R (Bürkner, 2016). Confidence was analysed with ordered logistic models, and accuracy was analyzed with Bernoulli models. In all cases, we used weakly informative priors (Normal(0,10)) for regression parameters, and performed 2000 iterations (burn-in period: 1000).

**Table S3.1:**
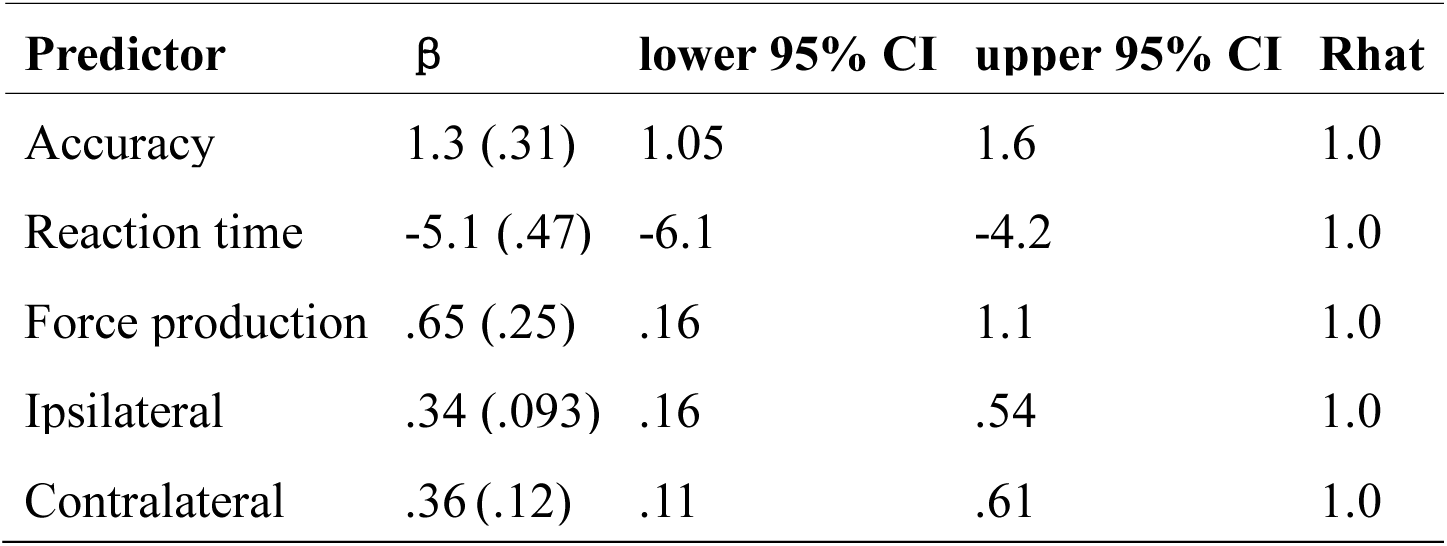
Predicting Confidence from Accuracy, Ispilateral and Contralateral partial activation, Reaction time and Force production. Predictors were coded as follows – Accuracy: error = 0,, correct = 1; Ipsilateral: absent = 0, present = 1; Contralateral: absent = 0, present = 1. Ordered logit model, implemented with brms package in R. Number of subjects: 19. Number of observations: 11471

| Predictor | $\beta$ | lower 95% CI | upper 95% CI | Rhat |
| --- | --- | --- | --- | --- |
| Accuracy | 1.3 (.31) | 1.05 | 1.6 | 1.0 |
| Reaction time | -5.1 (.47) | -6.1 | -4.2 | 1.0 |
| Force production | .65 (.25) | .16 | 1.1 | 1.0 |
| Ipsilateral | .34 (.093) | .16 | .54 | 1.0 |
| Contralateral | .36 (.12) | .11 | .61 | 1.0 |

**Table S3.2:** Predicting Confidence from Accuracy, Ispilateral and Contralateral partial activation, Reaction time and Force production. Predictors were coded as follows – Accuracy: error = 0, correct = 1; Ipsilateral: absent = 0, present = 1; Contralateral: absent = 0, present = 1. Ordered logit model, implemented with brms package in R.

| Predictor | $\beta$ | lower 95% CI | upper 95% CI | Rhat |
| --- | --- | --- | --- | --- |
| Accuracy | 1.2 (.16) | 1.04 | 1.7 | 1.0 |
| Reaction time | -5.4 (.52) | -6.5 | -4.4 | 1.0 |

| **Predictor** | **$\beta$** | **lower 95% CI** | **upper 95% CI** | **Rhat** |
| --- | --- | --- | --- | --- |
| Force production | .59 (.24) | .12 | 1.1 | 1.0 |
| Ipsilateral | .19 (.11) | -0.27 | .41 | 1.0 |
| Contralateral | .28 (.13) | .022 | .55 | 1.0 |
| Reaction time x Ipsilateral | 1.1 (.80) | .47 | 2.7 | 1.0 |
| Reaction time x Contralateral | 1.1 (.42) | .30 | 1.9 | 1.0 |

**Table S3.3:** Predicting Confidence from Accuracy, Ispilateral and Contralateral partial activation, Reaction time, Absolute Pre-Motor Time and Force production. Predictors were coded as follows – Accuracy: error = 0, correct = 1; Ipsilateral: absent = 0, present = 1; Contralateral: absent = 0, present = 1. Ordered logit, implemented with brms package in R. Number of subjects: 19. Number of observations: 11471

| **Predictor** | **$\beta$** | **lower 95% CI** | **upper 95% CI** | **Rhat** |
| --- | --- | --- | --- | --- |
| Accuracy | 1.3(.16) | 1.0 | 1.7 | 1.0 |
| Reaction time | -4.9 (.59) | -6.1 | -3.7 | 1.0 |
| Apmt | -.38 (.59) | -1.6 | .75 | 1.0 |
| Force production | .64 (.26) | .11 | 1.1 | 1.0 |
| Ipsilateral | .30 (.10) | .10 | .50 | 1.0 |
| Contralateral | .31 (.13) | .065 | .57 | 1.0 |

**Table S3.4:** Predicting Accuracy from Ispilateral and Contralateral partial activation, Reaction time, Absolute Pre-Motor Time and Force production. Predictors were coded as follows – Ipsilateral: absent = 0, present = 1; Contralateral: absent = 0, present = 1. Ordered logit, implemented with brms package in R.

| **Predictor** | **$\beta$** | **lower 95% CI** | **upper 95% CI** | **Rhat** |
| --- | --- | --- | --- | --- |
| Intercept | 1.7 (.25) | 1.2 | 2.2 | 1.0 |
| Reaction time | -2.3 (.45) | -3.2 | -1.4 | 1.0 |
| Force production | .14 (.21) | -.27 | .59 | 1.0 |
| Ipsilateral | -.095 (.17) | -.42 | .24 | 1.0 |
| Contralateral | .17 (.17) | -.14 | .54 | 1.0 |

**Table S3.5:** Predicting Accuracy from Ispilateral and Contralateral partial activation, Reaction time, Absolute Pre-Motor Time and Force production. Predictors were coded as follows –Ipsilateral: absent = 0, present = 1; Contralateral: absent = 0, present = 1. Ordered logit, implemented with brms package in R. Number of subjects: 19. Number of observations: 11471

| **Predictor** | **$\beta$** | **lower 95% CI** | **upper 95% CI** | **Rhat** |
| --- | --- | --- | --- | --- |
| Intercept | 1.7 (.27) | 1.2 | 2.3 | 1.0 |
| Reaction time | -2.4 (.48) | -3.3 | -1.4 | 1.0 |
| Force production | .12 (.21) | -.29 | .57 | 1.0 |
| Ipsilateral | -.38 (.19) | -.74 | .001 | 1.0 |
| Contralateral | .24 (.17) | -.074 | .59 | 1.0 |
| Reaction time x Ipsilateral | 1.3 (.56) | .25 | 2.4 | 1.0 |
| Reaction time x Contralateral | -.25 (.72) | -1.6 | 1.2 | 1.0 |

